# Using selective lung injury to improve murine models of spatially heterogeneous lung diseases

**DOI:** 10.1101/385922

**Authors:** Andrew J. Paris, Lei Guo, Ning Dai, Jeremy B. Katzen, Pryal N. Patel, G. Scott Worthen, Jacob S. Brenner

## Abstract

Many lung diseases, such as acute respiratory distress syndrome (ARDS), display significant regional heterogeneity, with patches of severely injured tissue adjacent to apparently healthy tissue. Current mouse models that aim to mimic ARDS generally produce diffuse injuries that cannot reproducibly generate ARDS’s regional heterogeneity. This deficiency prevents the evaluation of how well therapeutic agents reach the most injured regions, and precludes many regenerative medicine studies, since it is not possible to know which apparently healing regions suffered severe injury initially. Finally, these diffuse injury models must be mild to allow for survival, as their diffuse nature does not allow for residual healthy lung to keep an animal alive long enough for many drug and regenerative medicine studies. To solve all of these deficiencies of current animal models, we have created a simple and reproducible technique to selectively induce lung injury in specific areas of the lung. Our technique, catheter-in-catheter selective lung injury (CICSLI), involves guiding an inner catheter to a particular area of the lung and delivering an injurious agent mixed with nanoparticles (fluorescently and/or radioactively labeled) that can be used to track the location and extent of where the initial injury was, days later. Further, we demonstrate that CICSLI can produce a more severe injury than diffuse models, yet has much higher survival since CICSLI intentionally leaves undamaged lung regions. Collectively, these attributes of CICSLI will allow better study of how drugs act within heterogeneous lung pathologies and how regeneration occurs in severely damaged lung tissue, thereby aiding the development of new therapies for ARDS and other lung diseases.

## Introduction

Acute respiratory distress syndrome (ARDS) is characterized by patchy pulmonary infiltrates on chest x-ray, representing heterogeneously distributed inflammatory infiltrates and pulmonary edema[1–3]. While this heterogeneous distribution has been noted since the first descriptions of ARDS[4], it has been very difficult to study the consequences of regional heterogeneity of ARDS in experimental animal models. Animal models intended to model ARDS, called acute lung injury (ALI) models, such as intra-tracheal or instilled toxins (e.g., LPS) or intravenous toxins (e.g., oleic acid) produce a *diffuse* lung injury[5, 6]. When these models do produce heterogeneity[7], such as in the intra-nasal influenza model, the patchiness has very high inter-individual variability, largely limiting systematic study of the effects of patchy injury[8]. Further, because of the diffuse nature of such experimental ALI, the injuries must be kept mild to allow the animals to survive long enough for many analyses, such as for lung regeneration studies[7, 9].

These diffuse ALI models are particularly problematic for the primary two translational medicine fields aiming to ameliorate ARDS: development of therapeutics and regenerative medicine[10–15]. In the therapeutics space, the problem is that investigators must show that potential therapeutics for ARDS reach the patches of inflamed tissue (not just healthy lung regions), but diffuse ALI models do not allow such analyses because of their diffuse nature [15]. Thus with current diffuse ALI models we cannot determine the intra-pulmonary distribution of small molecule drugs, nano-scale drug delivery vehicles, inhaled drugs, or cell therapies. In the field of regenerative medicine, it is important to know whether a “healed” lung region was actually damaged, or if its healed appearance was because it was never injured in the first place. Since regenerative medicine requires analysis days to weeks after the injury, it is essential to be able to know the precise location and severity of the initial lung regions, even at long time points after injury[7, 9, 16]. Thus, for the development of ARDS therapeutics and post-ARDS lung regeneration treatments, the current diffuse ALI models leave room for optomiziation.

To solve those problems, we have created a very simple and reproducible system for creating and precisely tracking a heterogeneous and severe acute lung injury (ALI). In this system, termed catheter-in-catheter selective lung injury (CICSLI), mice are endo-tracheally intubated with a catheter[17], followed by insertion of a smaller catheter directed either into a single lung or single lobe, simply determined by the length of the smaller catheter. Into the smaller catheter a solution is instilled that contains an injurious insult (LPS, acid, etc) that has a low concentration of polymeric nanoparticles (labeled with fluorescence and/or radioactive moieties). This produces a severe single lung or single lobe ALI with excellent animal survival. Further, the nanoparticles allow tracing, several days later, of precisely where the injurious insult occurred, allowing fine determination of which regions received the initial injury, and correlating that with therapy distribution and lung regeneration assays.

## Materials and Methods

### Mice

All mice were housed in SPF conditions in an animal facility at the Children’s Hospital of Philadelphia. All mouse protocols were approved by the IACUC at the Children’s Hospital of Philadelphia. WT C57BL/6J mice (strain 000664 from the Jackson Laboratory, Bar Harbor, ME) aged 8-10 weeks were used for experiments. Both male and female mice were used in equal proportions.

### Injury Model

Sedated mice were intubated using a 20G angiocatheter (BD catalog #381434) using a previously described technique[17]. The mice were then placed in the right lateral recumbent position at a polytehelene 10 (PE-10) catheter (BD catalog #427400) was directed into the right main stem bronchus. Injury was induced by instilling 2 μL/g of osmotically balanced 0.1N HCl into the right lung through the PE-10 catheter.

### Instillation of LPS

Lipopolysaccharide B4 (Sigma catalog # L2630) was selectively instilled into mouse airways, as described in sections above. Each mouse was instilled with 1 mg/kg of LPS.

### Nanoparticle-based tracking of the location of instillate into airways

Polymeric nanoparticles (NPs) were purchased from Bangs Laboratories, Inc. For tracking where the NPs localized by immunofluorescence days after acid instillation, we used NPs that were 1000 nanometer (nm) in diameter and composed of polystyrene with the fluorphore FlashRed (similar spectrum to Cy5) covalently attached (Bangs Labs Catalog #FSFR004). For radiotracing where the instillate localized, we used 200-nm polystyrene conjugated to FlashRed and with surface carboxylate groups (Catalog #FSFR002). The surface carboxylation allowed for conjugation to radiolabeled proteins, as described below.

### Radiolabeling nanoparticles

Rat IgG (ThermoFisher catalog #31933) was labeled with I-125 via Pierce Iodination Beads (ThermoFisher catalog # 28665). We then conjugated the IgG to nanoparticles using our published protocol[15]. In brief, 100 μL of polystyrene NPs were buffer exchanged Zeba™ Spin Desalting Columns, 7K MWCO, 0.5 mL (ThermoFisher catalog # 89882), exchanging for 50 mM MES buffer at pH 5.2, finally putting the buffer-exchanged beads into 1.5 mL Eppendorf tubes. Next N-Hydroxysulfosuccinimide (“sulfo-NHS”; Sigma catalog #56485) was added to a final concentration of 0.275 mg/ml and incubate for 3 minutes at room temperature (RT) Next N-(3-Dimethylaminopropyl)-N′-ethylcarbodiimide hydrochloride (“EDAC”; Sigma catalog # E7750) was added to a final concentration of 0.1 mg/ml and incubated for 10-15 minutes at RT. Next 114 ug of I-125-labeled rat IgG was added (giving 200 antibody molecules per bead) and incubated for 2 to 4 hours at RT on a vortex/shaker at low speed. 1 mL MES buffer was added to dilute free antibody, followed by centrifuge at 12,000g x 3 min to pellet the IgG-conjugated NPs. The IgG-NP pellet was resuspended in 200 uL of PBS + 0.05% bovine serum albumin (BSA) buffer. Immediately before use, the NPs were sonicated with a probe / tip sonicator for three 3-second pulses at 30% maximum power.

### Radiolabeled albumin tracing

BSA was I-125-labeled as described for IgG above. Mice were given selective instillation of LPS, followed 20 hours later by intravenous injection of 1 x 10^6^ counts per minute (cpm) of I-125-BSA, followed by sacrifice 4 hours later. The right ventricle was then flushed with 10 mL of PBS to flush out the pulmonary vasculature of residual blood. The lobes of the lungs were then individually removed from chest cavity and measured for I-125 levels in a Perkin Elmer Wizard-2 gamma counter.

### Evans Blue determination of capillary leak

Mice were given lobar LPS. Twenty four hours later, mice were injected IV with Evans Blue at 30 mg/kg, followed 2 hours later by perfusing the right ventricle with 10 mL of PBS + 10mM EDTA. The lung lobes were individually dissected from the chest and a photograh was taken of them.

### CT scanning of mice

Mice were given lobar LPS. Twenty four hours later, under general anesthesia a tracheostomy was created and a 20 gauge peripheral IV catheter was placed into the tracheostomy. The mouse was then immediately sacrificed via overdose of ketamine. Before the development of post-mortem atelectasis, the tracheostomy tube was removed while a ligature was cinched around the trachea, thus ensuring the lungs remained filled with air. Immediately after sacrifice, the mouse’s body was put into a small animal microCT scanner made by ImTek, housed at the University of Pennsylvania’s Small Animal Imaging Facility.

## Results

### A simple and traceable method for selectively injuring murine lungs

We developed a straightforward and minimally invasive technique for instilling liquid into a mouse’s lung. The procedure starts with intubating the mouse using a previously established method of orotracheal intubation using a 20g angiocatheter[17]. Once the angiocatheter has been advanced into the mouse’s trachea, we then insert polyethylene 10 tubing (54 mm) attached to a 1mL syringe via a 30G needle (Fig 1A). For the purpose of demonstrating the method in the left lung we used a 2uL per gram of methylene blue followed by 200ul of air. The mouse was placed on in the left lateral recumbent position before slowly instilling the methylene blue. Afterwards, the mouse was placed in the supine position and the lungs were extracted and photographed to demonstrate the that this method can achieve selective instillation of a liquid into a living mouse (Fig 1B).

**Fig 1.**
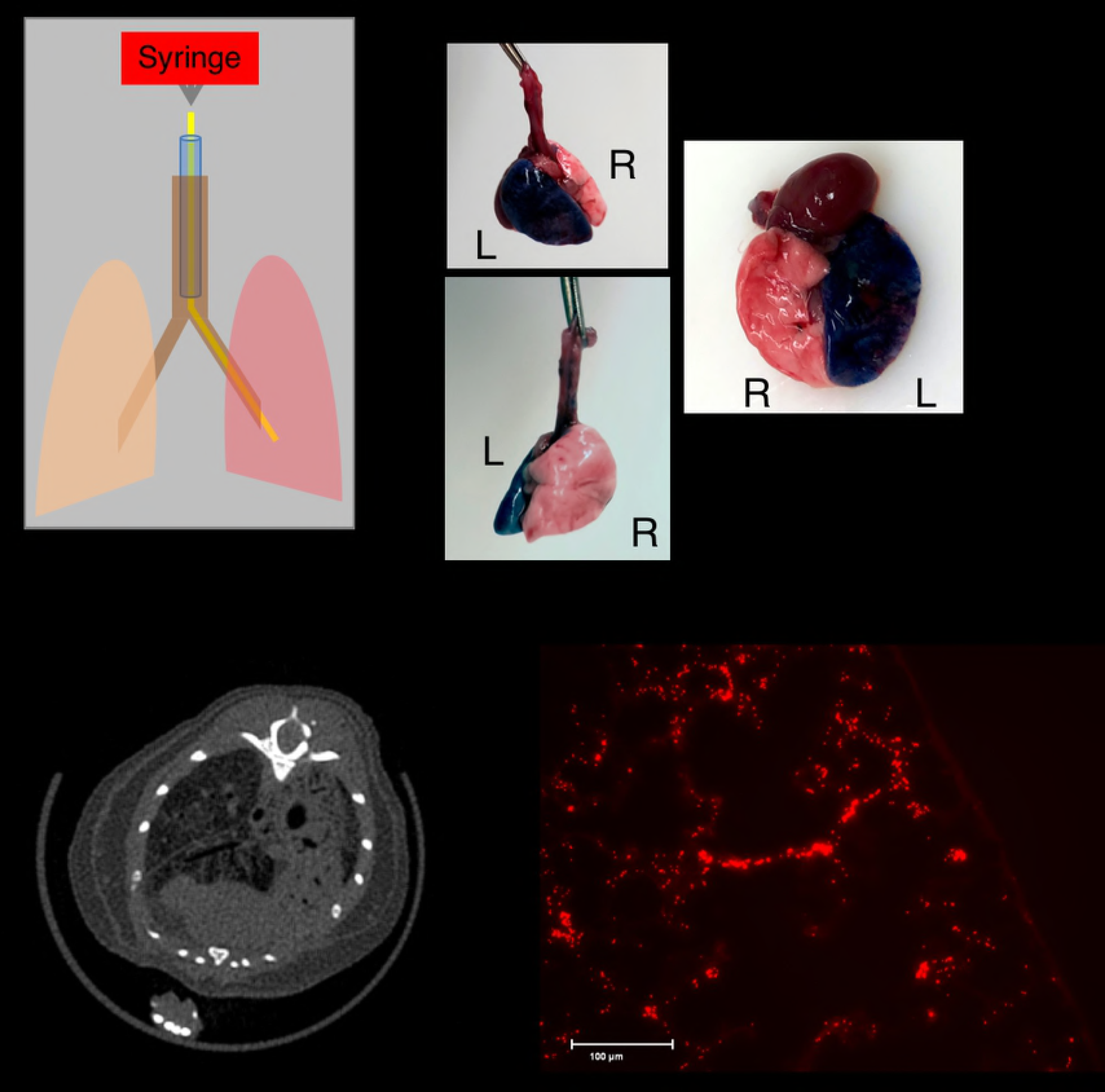
Unilateral lung injury, created by instilling acid into just one lung. A) Procedure for single lung acute injury. The mouse was sedated, followed by orotracheal intubation with a 20G angiocatheter (Blue) that terminates in the trachea (brown). The mouse is then placed in the left lateral recumbent position and a 51mm PE10 catheter (yellow) is advanced into the inferior lung (the left lung in this example). A syringe is attached to the catheter using a 0.5 inch 30G needle (not shown) and a specified amount of liquid is selectively delivered to the inferior lung (red). B) Healthy mice underwent selective instillation of methylene blue as detailed in A. The mouse was euthenized 5 minutes after instillation. Selective injury of the left lung was verified visually. C) Selective instillation of 0.1N osmotically balanced hydrochloric acid into the right lung was performed 24 hours before euthanizing the mice and subejecting them to microCT scans. CT imaging shows parenchymal injury exclusively on the right side. D) Mice underwent single lung instillation of acid containing 1 μM FlashRed-labeled polystyrene beads, followed by euthanasia 72 hours later. Frozen sections of embedded injured lung tissue demonstrated persistence of beads at 72 hours post-injury.

To demonstrate that this method can be used to induce selective lung injury we repeated the procedure outlined in Fig 1A but instead placed the mouse in the right lateral recumbent position in order to guide the PE 10 catheter into the right lung and instilled 2.5ul per gram of 0.1N hydrochloric acid into the mouse’s right lung. A CT scan performed one day later showed selective injury of the right lung, indicating that we can safely induce a radiographically apparent injury without causing the mouse to die (Fig 1C). Because this method is ideal for studying lung regeneration we wanted to find a way to distinguish where injury occurred. Ordinarily, it may not be possible to distinguish between uninjured and properly regenerated lung tissue, therefore we sought to mark where injury had occurred by including 1uM of innocuous FlashRed-labeled polystyrene beads. Frozen section of the injured right lung demonstrate that the beads are readily apparent even when delivered in a solution containing 1M hydrochloric acid (Fig 1D). No beads were visible in the contralateral, uninjured, lung (data not shown).

### Selective lung injury paradoxically increases histological injury and decreases mortality

To ascertain the relative difference in the relative differences in tissue injury between unilateral and diffuse lung injury, we instilled 2.5ul/g of 1N hydrochloric acid into the lungs of C57BL/6 mice using our selective method (Fig 1) or intratracheally, meaning without using the PE10 catheter. We observed a relatively more severe lung injury in the selective group with a markedly increased cellular infiltrate and hyaline membrane formation. The mice that received bilateral lung injury had less cellular infiltrate (Fig 2A). Although intratracheal administration produced a more bland-appearing injury, the survival at twenty-four hours was significantly lower in the mice that had bilateral lung injury relative to those that had unilateral injury (Fig 2B). These data reveal a paradox in which selectively injuring lungs increases histological injury while decreasing mortality.

**Fig 2.**
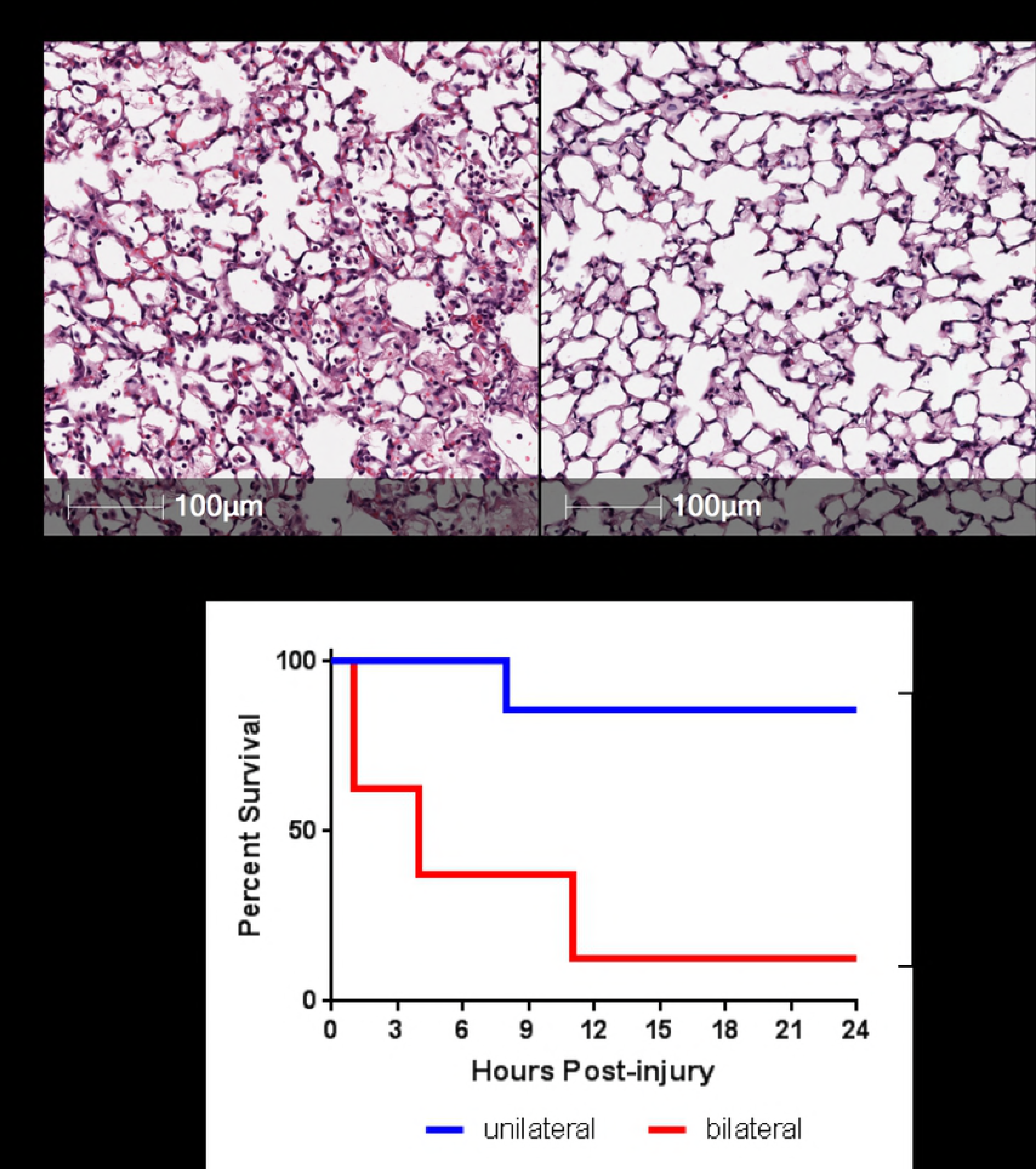
Unilateral lung injury increases injury while improving survival. A) H&E staining of lung sections taken from C57BL/6 mice 24 hours after undergoing unilateral or bilateral lung injury demonstrates significantly increased accumulation of cellular infiltration and protein-rich edema fluid when compared to the lung that underwent bilateral (intratracheal) lung injury. B) We subjected mice to right unilateral (n=7) and bilateral (n=8) lung injury. Mice were evaluated on an hourly basis and were euthanized if a blinded observer determined that the mouse was obtunded. Differences in mortality were significant, *p* = 0.0055, when compared using a log-rank (Mantel-Cox) test.

### Selectively injuring a single lobe is possible and remains isolated

Data shown in Figures 1 and 2 demonstrate that instilling agents through a catheter that is longer than the endotracheal tube can help selectively injure the left (Fig 1A and B) or right (Fig 1C) lungs. Based on these data we hypothesized that using a longer catheter and smaller volume of fluid could selectively injure a single lobe of the right lung. To test this hypothesis, we repeated the experimental design shown in Figure 1 but used a slightly longer catheter – 59mm PE 10 catheter (Fig.3A). Using Evan's blue dye, we show that the longer catheter is able to selectively instill fluid into the RUL (Fig 3B). The immediate specificity of selectively instilling lipopolysaccharide (LPS) into the right upper lobe was assayed by co-instilling I-125-labeed-IgG that was bound to 100nm polystyrene beads along with the LPS. We observed that 97% of the radiolabeled albumin was detected in the RUL, quantifying the selectiveness of this approach (Fig 3C). To determine if the initial instillation remained isolated in the right upper lobe we repeated the same test with I-125-labeled IgG co-administered with LPS and found that 92% of the isotope was detected in the right upper lobe 24 hours after selectively instilling LPS in the right upper lobe, suggesting that the selective lung injury did not spread to other lobes.

**Fig 3.**
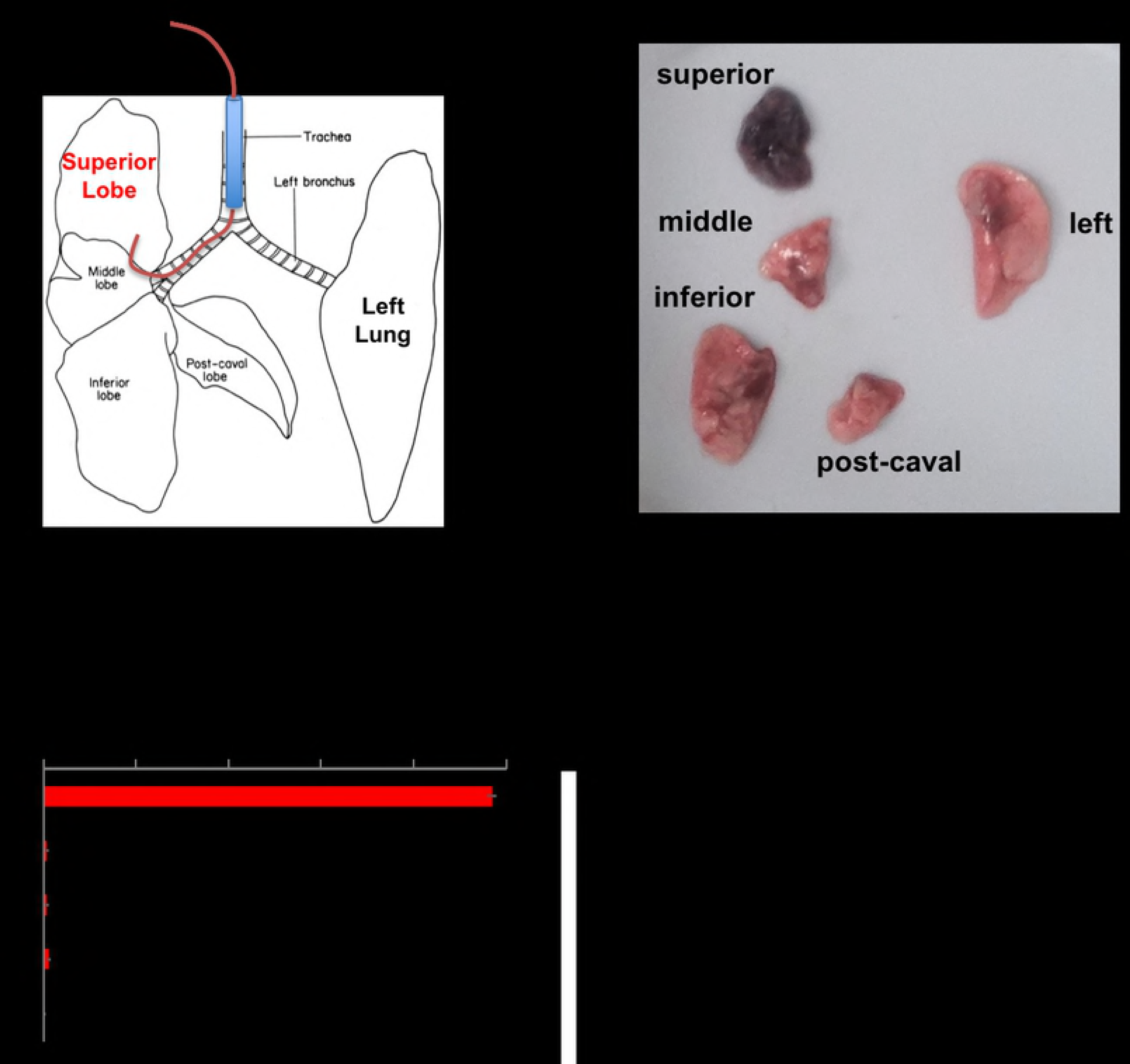
Single-lobe injury created by a longer instillation catheter: A) Mice were treated as in Figure 1, but instead a 59mm PE10 catheter was used and the mice are kept in left lateral recumbent for 20 minutes after instillation. B) Mice were instilled as in A, with the instillate containing 1mg/kg of LPS, 24 hours later, Evans Blue dye was injected followed by sacrifice 2 hours later, with visible photography of the lobes. C) Mice were treated as in C, but into the LPS mixture was included I-125-labeled-IgG coated onto 100 nanometer diameter polystyrene beads. Immediately after instillation, the mice were sacrificed, followed by measurement of each lobe in a gamma counter. Each data point represents mean ± s.e.m (n=3). * p<0.0001, one-way ANOVA.

### Selective lung injury changes regional lung physiology physiology

Hematoxylin and eosin (H&E) staining of a lung following lung injury revealed a spectrum of tissue damage within the injured lung, which we attribute to uneven distribution of the LPS within the injured lobe. Importantly, we did not observe any areas of uninjured tissue within the injured lobe (Fig 4A). Although selective injury was able to induce lobe-specific tissue damage (Figs 1C, 4A) we were uncertain if a seminal physiologic response to injury was also regional or if the severe injury would induce systemic changes in all lobes. To assay for vascular leak, we injected I-125-labeled albumin via retroorbital injection 24 hours after inducing selective lung injury. When compared to uninjured mice we found that only the right upper lobe had a significantly increased amount of radiolabeled albumin (Fig 4B). These data suggest that selective injury can recapitulated lobe-specific tissue damage and physiologic change.

**Fig 4.**
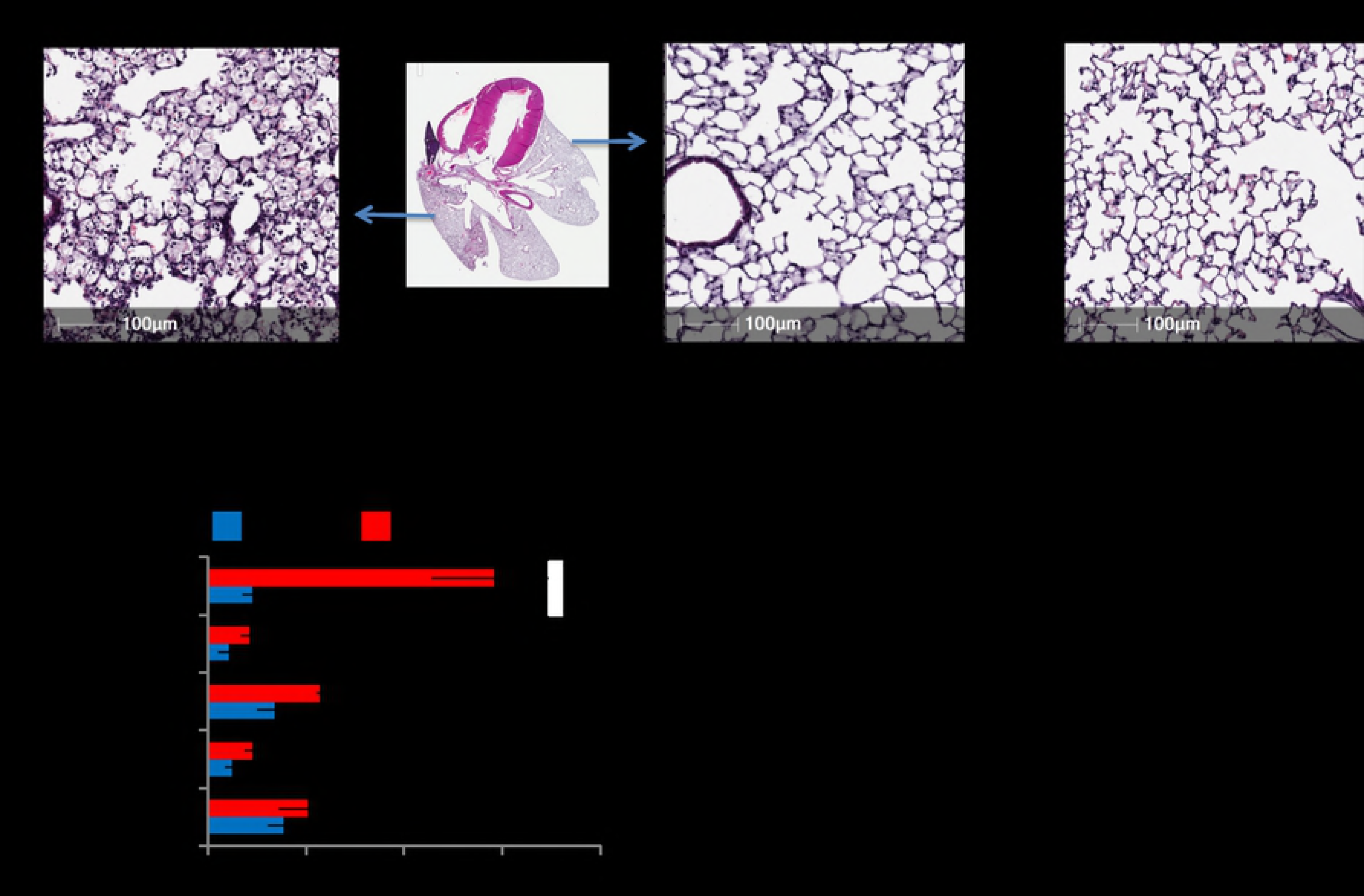
In single-lobe lung injury, some pathological phenotypes are restricted to injured lobes. A) Mice underwent single-lobe LPS instillation as in Figure 3, followed by sacrifice 24 hours later, and then H&E staining of the lungs. Comparison is shown to a naïve mouse. B) Mice underwent single-lobe LPS instillation, followed by injection of I-125-labeled albumin 24 hours later, followed by sacrifice, perfusion of the pulmonary arteries with 5 mL of PBS, and then gamma counting of the lobes. Note that the lobe into which LPS was instilled (the superior) has a greater capillary leak than the other lobes. Each data point represents mean ± s.e.m (n=3). * p<0.0001, one-way ANOVA, followed by pairwise comparison for each lobe, comparing naïve vs single lobe LPS, with all lobes having non-significant differences except the superior lobe.

## Discussion

Developing and testing new therapeutics for lung diseases starts with using outstanding pre-clinical models that recapitulate key aspects of human lung disease[15, 18]. Intratracheal administration of injurious agents or inducing non-specific systemic inflammation either create bland injury that falls short of the severe injury that often produces the greatest mortality or it creates such a severe injury that studying long-term regeneration is not possible. Furthermore, inducing regional physiologic changes can help us understand how drugs will be distributed in function in a patient with a mix of injured and healthy tissues[15], which is the case in most human diseases[3, 19–21].

In this paper we describe a simple and reproducible technique to selectively injure specific parts of the lung. By combining a catheter-within-a-catheter technique and positioning a mouse in the right or left lateral recumbent position we can easily direct a lung injury to the right or left lung. Further lengthening the inner catheter facilitates selective injury of the right upper lobe. Inducing injury in this way is the major innovation of our model because of its ability to produce a histologically severe yet simultaneously more survivable injury. This has allowed us to study how regeneration following a tissue injury that would otherwise be unsurvivable[22].

We considered that using these models to study lung regeneration would be challenging as newly regenerated lung tissue would appear histologically identical to the uninjured lung. Although lineage tracing[23] of alveolar epithelial progenitors could be used to overcome this challenge, there is no way label all progenitor populations[7, 9, 16, 24, 25], meaning that regenerated lung tissue could still resemble uninjured lung even when using lineage tracing. Labeling the injured area is thus an essential second innovation that we are reporting. We show here that co-administration of innocuous fluorescently-labeled beads or radiolabeled albumin will persist as validated markers of where injury occurred.

We have also shown that our selective lung injury model is useful for inducing selective changes in lung permeability. This third innovation recapitulates key heterogeneity that occurs in our patients[2, 3, 26] and has important implications for evaluating candidate drugs. Indeed, we have already shown regional changes in drug delivery which may be useful in understanding how to selectively deliver drugs to injured lungs while avoiding normal tissue.

Although other groups have published reports utilizing selective lung injury[27, 28] we believe that this is the first systematic approach to delineating the methodology and interrogating the physiology though the lens of developing better pre-clinical mouse models of lung disease. Further work is needed to determine best-practices for histologically scoring heterogenous lung injuries and assaying lung function. Furthermore, technological advances, such as the recently described mouse bronchoscope[29], may facilitate repeat instillations and sequential sampling. Nevertheless, this report details a simple, low-cost, and low-labor method that can be easly employed that can dramatically improve the study of ARDS.

Notably, the CICSLI technique can be used to prepare not just ARDS models, but perhaps other heterogeneous lung diseases. Numerous other lung diseases are modeled by intra-tracheal instillations: pneumonia with bacteria[30], idiopathic pulmonary fibrosis with bleomycin[31], and asthma with airway irritants[32]. Each of these human diseases is very spatially heterogeneous and so would benefit from CICSLI. Additionally, CICSLI could be done with new agents, such as transfection with gene therapy and CRISPR reagents, to create new pathology models.

In summary, CICSLI provides a simple, easy, quick and inexpensive method to improve key features of pre-clinical lung disease models, which should aid the development of new drugs and regenerative medicine therapies.

## Acknowldegments

This work was funded through the generous support of the Univesrty of Pennsylvania Measey Physician Scientist Fellowship Alward (AJP), the Parker B. Francis Fellowship Program (AJP), and multiple grants form the NIH – K08 HL136698 (AJP), K08 HL138269 (JSB), R01 AI099479 (GSW) and UG3 TR002198 (GSW).

## References

1. Bernard G. Acute Lung Failure - Our Evolving Understanding of ARDS. The New England journal of medicine. 2017;377(6):507-9. Epub 2017/08/10. doi: 10.1056/NEJMp1706595. PubMed PMID: 28792872.

2. Force ADT, Ranieri VM, Rubenfeld GD, Thompson BT, Ferguson ND, Caldwell E, et al. Acute respiratory distress syndrome: the Berlin Definition. Jama. 2012;307(23):2526-33. doi: 10.1001/jama.2012.5669. PubMed PMID: 22797452.

3. Thompson BT, Chambers RC, Liu KD. Acute Respiratory Distress Syndrome. The New England journal of medicine. 2017;377(19):1904-5. Epub 2017/11/09. doi: 10.1056/NEJMc1711824. PubMed PMID: 29117492.

4. Ashbaugh DG, Bigelow DB, Petty TL, Levine BE. Acute respiratory distress in adults. Lancet. 1967;2(7511):319-23. Epub 1967/08/12. PubMed PMID: 4143721.

5. Matute-Bello G, Downey G, Moore BB, Groshong SD, Matthay MA, Slutsky AS, et al. An official American Thoracic Society workshop report: features and measurements of experimental acute lung injury in animals. Am J Respir Cell Mol Biol. 2011;44(5):725-38. doi: 10.1165/rcmb.2009-0210ST. PubMed PMID: 21531958.

6. Matute-Bello G, Frevert CW, Martin TR. Animal models of acute lung injury. American journal of physiology Lung cellular and molecular physiology. 2008;295(3):L379-99. doi: 10.1152/ajplung.00010.2008. PubMed PMID: 18621912; PubMed Central PMCID: PMC2536793.

7. Zacharias WJ, Frank DB, Zepp JA, Morley MP, Alkhaleel FA, Kong J, et al. Regeneration of the lung alveolus by an evolutionarily conserved epithelial progenitor. Nature. 2018;555(7695):251-5. Epub 2018/03/01. doi: 10.1038/nature25786. PubMed PMID: 29489752; PubMed Central PMCID: PMCPMC6020060.

8. Morales-Nebreda L, Chi M, Lecuona E, Chandel NS, Dada LA, Ridge K, et al. Intratracheal administration of influenza virus is superior to intranasal administration as a model of acute lung injury. J Virol Methods. 2014;209:116-20. Epub 2014/09/23. doi: 10.1016/j.jviromet.2014.09.004. PubMed PMID: 25239366; PubMed Central PMCID: PMCPMC4315182.

9. Zuo W, Zhang T, Wu DZ, Guan SP, Liew AA, Yamamoto Y, et al. p63(+)Krt5(+) distal airway stem cells are essential for lung regeneration. Nature. 2015;517(7536):616-20. doi: 10.1038/nature13903. PubMed PMID: 25383540.

10. Chapman HA. Toward lung regeneration. The New England journal of medicine. 2011;364(19):1867-8. doi: 10.1056/NEJMe1101800. PubMed PMID: 21561353.

11. Matthay MA, Wiener-Kronish JP. Intact epithelial barrier function is critical for the resolution of alveolar edema in humans. The American review of respiratory disease. 1990;142(6 Pt 1):1250-7. doi: 10.1164/ajrccm/142.6_Pt_1.1250. PubMed PMID: 2252240.

12. Cross LJ, O'Kane CM, McDowell C, Elborn JJ, Matthay MA, McAuley DF. Keratinocyte growth factor in acute lung injury to reduce pulmonary dysfunction--a randomised placebo-controlled trial (KARE): study protocol. Trials. 2013;14:51. doi: 10.1186/1745-6215-14-51. PubMed PMID: 23419093; PubMed Central PMCID: PMCPMC3620926.

13. Matthay MA, Liu KD. New Strategies for Effective Therapeutics in Critically Ill Patients. Jama. 2016;315(8):747-8. doi: 10.1001/jama.2016.0661. PubMed PMID: 26903328.

14. Standiford TJ, Ward PA. Therapeutic targeting of acute lung injury and acute respiratory distress syndrome. Transl Res. 2016;167(1):183-91. doi: 10.1016/j.trsl.2015.04.015. PubMed PMID: 26003524; PubMed Central PMCID: PMCPMC4635065.

15. Brenner JS, Bhamidipati K, Glassman P, Ramakrishnan N, Jiang D, Paris AJ, et al. Mechanisms that determine nanocarrier targeting to healthy versus inflamed lung regions. Nanomedicine. 2017. doi: 10.1016/j.nano.2016.12.019. PubMed PMID: 28065731.

16. Vaughan AE, Brumwell AN, Xi Y, Gotts JE, Brownfield DG, Treutlein B, et al. Lineage-negative progenitors mobilize to regenerate lung epithelium after major injury. Nature. 2015;517(7536):621-5. doi: 10.1038/nature14112. PubMed PMID: 25533958; PubMed Central PMCID: PMC4312207.

17. Das S, MacDonald K, Chang HY, Mitzner W. A simple method of mouse lung intubation. J Vis Exp. 2013;(73):e50318. doi: 10.3791/50318. PubMed PMID: 23542122; PubMed Central PMCID: PMCPMC3639692.

18. Vlahos R, Bozinovski S. Recent advances in pre-clinical mouse models of COPD. Clin Sci (Lond). 2014;126(4):253-65. Epub 2013/10/23. doi: 10.1042/CS20130182. PubMed PMID: 24144354; PubMed Central PMCID: PMCPMC3878607.

19. Bellani G, Laffey JG, Pham T, Fan E, Brochard L, Esteban A, et al. Epidemiology, Patterns of Care, and Mortality for Patients With Acute Respiratory Distress Syndrome in Intensive Care Units in 50 Countries. Jama. 2016;315(8):788-800. doi: 10.1001/jama.2016.0291. PubMed PMID: 26903337.

20. Lederer DJ, Martinez FJ. Idiopathic Pulmonary Fibrosis. The New England journal of medicine. 2018;378(19):1811-23. Epub 2018/05/10. doi: 10.1056/NEJMra1705751. PubMed PMID: 29742380.

21. Musher DM, Thorner AR. Community-acquired pneumonia. The New England journal of medicine. 2014;371(17):1619-28. Epub 2014/10/23. doi: 10.1056/NEJMra1312885. PubMed PMID: 25337751.

22. Paris AJ, Liu Y, Mei J, Dai N, Guo L, Spruce L, et al. Neutrophils promote alveolar epithelial regeneration by enhancing type II pneumocyte proliferation in a model of acid-induced acute lung injury. American journal of physiology Lung cellular and molecular physiology. 2016:ajplung 00327 2016. doi: 10.1152/ajplung.00327.2016. PubMed PMID: 27694472.

23. Kretzschmar K, Watt FM. Lineage tracing. Cell. 2012;148(1-2):33-45. doi: 10.1016/j.cell.2012.01.002. PubMed PMID: 22265400.

24. Barkauskas CE, Cronce MJ, Rackley CR, Bowie EJ, Keene DR, Stripp BR, et al. Type 2 alveolar cells are stem cells in adult lung. The Journal of clinical investigation. 2013;123(7):3025-36. doi: 10.1172/JCI68782. PubMed PMID: 23921127; PubMed Central PMCID: PMCPMC3696553.

25. Ray S, Chiba N, Yao C, Guan X, McConnell AM, Brockway B, et al. Rare SOX2+ Airway Progenitor Cells Generate KRT5+ Cells that Repopulate Damaged Alveolar Parenchyma following Influenza Virus Infection. Stem Cell Reports. 2016;7(5):817-25. doi: 10.1016/j.stemcr.2016.09.010. PubMed PMID: 27773701; PubMed Central PMCID: PMCPMC5106521.

26. Guerin C, Reignier J, Richard JC, Beuret P, Gacouin A, Boulain T, et al. Prone positioning in severe acute respiratory distress syndrome. The New England journal of medicine. 2013;368(23):2159-68. doi: 10.1056/NEJMoa1214103. PubMed PMID: 23688302.

27. Amigoni M, Bellani G, Scanziani M, Masson S, Bertoli E, Radaelli E, et al. Lung injury and recovery in a murine model of unilateral acid aspiration: functional, biochemical, and morphologic characterization. Anesthesiology. 2008;108(6):1037-46. doi: 10.1097/ALN.0b013e318173f64f. PubMed PMID: 18497604.

28. Fukunaga K, Kohli P, Bonnans C, Fredenburgh LE, Levy BD. Cyclooxygenase 2 plays a pivotal role in the resolution of acute lung injury. Journal of immunology. 2005;174(8):5033-9. Epub 2005/04/09. PubMed PMID: 15814734.

29. Dames C, Akyuz L, Reppe K, Tabeling C, Dietert K, Kershaw O, et al. Miniaturized bronchoscopy enables unilateral investigation, application, and sampling in mice. Am J Respir Cell Mol Biol. 2014;51(6):730-7. Epub 2014/06/25. doi: 10.1165/rcmb.2014-0052MA. PubMed PMID: 24960575.

30. Nuermberger E. Murine models of pneumococcal pneumonia and their applicability to the study of tissue-directed antimicrobials. Pharmacotherapy. 2005;25(12 Pt 2):134S-9S. Epub 2005/11/25. doi: 10.1592/phco.2005.25.12part2.134S. PubMed PMID: 16305283.

31. Mouratis MA, Aidinis V. Modeling pulmonary fibrosis with bleomycin. Curr Opin Pulm Med. 2011;17(5):355-61. Epub 2011/08/13. doi: 10.1097/MCP.0b013e328349ac2b. PubMed PMID: 21832918.

32. Kips JC, Anderson GP, Fredberg JJ, Herz U, Inman MD, Jordana M, et al. Murine models of asthma. Eur Respir J. 2003;22(2):374-82. Epub 2003/09/04. PubMed PMID: 12952276.

